# IER3-N-cadherin axis contributes to apoptosis resistance in dermal fibroblasts under profibrotic conditions

**DOI:** 10.64898/2026.10.31.763371

**Authors:** Ryuzo Ichimura, Hibari Nakajima, Ayaka Sugimori, Hayakazu Sumida

**Author notes:** **Corresponding author** Hayakazu Sumida Department of Dermatology, Faculty of Medicine, The University of Tokyo, 7-3-1, Hongo, Bunkyo-ku, Tokyo, 113-8655, Japan.

## Abstract

**Background:** Persistent activation and resistance to apoptosis of dermal fibroblasts contribute to pathological skin fibrosis. Immediate early response 3 (IER3) is a stress-responsive protein involved in cell survival, but its role in cutaneous fibrosis remains unclear. We investigated the functional significance of IER3 in dermal fibroblasts and its potential contribution to fibrosis.

**Methods:** IER3 expression was examined in skin samples from patients with systemic sclerosis, keloids, and hypertrophic scars. Primary human dermal fibroblasts were stimulated with transforming growth factor-β1 (TGF-β1). RNA sequencing, siRNA-mediated gene silencing, quantitative PCR, and flow cytometric apoptosis assays were performed to investigate the downstream mechanisms and functional consequences of IER3 expression.

**Results:** IER3 expression was increased in dermal fibroblasts within fibrotic skin lesions and was induced by TGF-β1 independently of SMAD3 and MEK/ERK signaling. Transcriptomic analysis identified *CDH2*, encoding N-cadherin, as a downstream target of IER3. Moreover, TGF-β1 induced *CDH2* expression in a SMAD3 phosphorylation-dependent manner. *IER3* knockdown attenuated TGF-β1-induced *CDH2* expression, whereas *CDH2* knockdown did not affect *IER3* expression. Furthermore, *CDH2* knockdown enhanced staurosporine-induced apoptosis in TGF-β1-stimulated fibroblasts, accompanied by a shift in apoptosis-related gene expression toward a pro-apoptotic profile.

**Conclusion:** Our findings identify a previously unrecognized IER3-N-cadherin regulatory axis that may promote dermal fibroblast survival by suppressing apoptosis under profibrotic conditions. Targeting this pathway may represent a novel therapeutic strategy for persistent skin fibrosis.

## INTRODUCTION

Fibrosis is a pathological process characterized by excessive accumulation of extracellular matrix (ECM), including collagen, that can impair physiological function. Under physiological conditions, fibrosis is needed for tissue repair with a regulated sequence of chronic inflammation and fibroblast activation. Fibroblasts play a central role in these processes by producing and remodeling ECM components [1]. Upon activation by profibrotic stimuli, fibroblasts acquire a myofibroblast phenotype characterized by expression of α-smooth muscle actin (α-SMA) and enhanced production of ECM proteins [1]. During normal wound healing, activated fibroblasts and myofibroblasts are eliminated or return to a quiescent state as tissue repair progresses [1, 2]. In contrast, persistent activation and resistance to apoptosis of fibroblasts and myofibroblasts, with sustained and excessive production of profibrotic mediators such as transforming growth factor-β (TGF-β), can promote excessive ECM accumulation and progressive tissue fibrosis [1, 3].

Tissue remodeling with fibrosis is a major pathological feature of various chronic diseases affecting multiple organs, including the lung, liver, kidney, heart, and skin [4]. Importantly, ECM deposition in fibrosis can induce irreversible structural/functional alterations and disruption of normal tissue architecture. Severe irreversible fibrosis is often resistant to currently available therapeutic interventions, highlighting the unmet need for a better understanding of the molecular mechanisms underlying the persistence of fibrotic lesions and the development of novel antifibrotic strategies [5]. Especially in the skin, a variety of fibrotic disorders are characterized by excessive collagen deposition and activation of dermal fibroblasts/myofibroblasts. These conditions include pathological scar formation such as hypertrophic scars and keloids, which remain clinically challenging because of their persistent fibrotic changes, recurrence after treatment, and limited therapeutic options [6]. Despite ongoing efforts to overcome this challenge, the mechanisms responsible for sustained fibroblast activation remain incompletely understood. Regarding systemic fibrotic diseases, systemic sclerosis (SSc) is a prototypic and heterogeneous autoimmune disorder characterized by fibrosis of the skin and certain organs. SSc is driven by a combination of immune dysregulation, vascular abnormalities, and aberrant fibroblast activation, leading to excessive ECM accumulation and resulting in tissue fibrosis and functional impairment [7]. Although several immune-targeting or anti-fibrotic therapies have been recently introduced for the management of fibrotic diseases, effective therapeutic strategies directly targeting established fibrosis remain limited [8]. Thus, therapeutic strategies aimed at eliminating persistent profibrotic fibroblasts or restoring their susceptibility to apoptosis may offer a promising approach to limiting fibrosis and promoting the resolution of established fibrotic lesions.

Oxidative stress has been implicated as a critical contributor to the initiation and progression of various fibrotic diseases. Reactive oxygen species (ROS) serve as important signaling molecules that regulate cellular responses. Under physiological conditions, ROS production is tightly controlled by antioxidant systems; however, excessive or sustained ROS accumulation can induce oxidative damage and promotes profibrotic signaling pathways [9]. Increased oxidative stress has been documented in several fibrotic disorders, including SSc, lung fibrosis, and liver fibrosis, suggesting that redox imbalance may be related to fibrotic progression across multiple organs [10, 11]. In particular, ROS can drive fibrosis by activating fibroblasts and enhancing TGF-β signaling, leading to myofibroblast differentiation and excessive ECM deposition [12, 13]. Despite growing evidence linking oxidative stress to fibrosis, the molecular mechanisms linking stress responses to fibrosis remain incompletely understood. Immediate early response 3 (IER3), also known as IEX-1, is a stress-responsive protein implicated in the regulation of cellular survival and oxidative stress responses [14, 15]. However, its functional role in dermal fibroblasts and its potential contribution to cutaneous fibrosis remain unclear. Therefore, in this study, we aimed to investigate the role of IER3 in dermal fibroblast function and its potential involvement in the pathogenesis of fibrotic skin diseases.

## MATERIALS AND METHODS

### Human skin samples

Skin samples were obtained from lesional skin of five patients with systemic sclerosis (SSc), three patients with keloids, and three patients with hypertrophic scars. As controls, normal skin samples were obtained from five healthy individuals. All patients with SSc fulfilled the 2013 American College of Rheumatology/European League Against Rheumatism (ACR/EULAR) classification criteria for SSc [16]. All samples were collected from individuals who visited the Department of Dermatology, The University of Tokyo Hospital. All experimental protocols involving human samples were approved by the Ethics Committee of The University of Tokyo (approval nos. 0695, 10819, 2023051G, and 2024385G). Written informed consent was obtained from all participants in accordance with the principles of the Declaration of Helsinki.

### Cell culture and stimulation

Primary human dermal fibroblasts were isolated from clinical skin samples as previously described [17]. Briefly, skin samples were cut into small pieces (several millimeters in size), placed on culture dishes, and maintained in minimum essential medium (MEM) supplemented with 10% fetal bovine serum (FBS). After several days of culture, fibroblasts migrated out from the tissue explants and began to proliferate. Primary human dermal fibroblasts were cultured in MEM supplemented with 10% fetal bovine serum (FBS; BioWest, Nuaillé, France) and antibiotics (penicillin/streptomycin; Nacalai Tesque, Kyoto, Japan). Cells were maintained as monolayer cultures at 37°C in a humidified atmosphere containing 5% CO₂. Fibroblasts between passages 2 and 5 were used for all experiments. Six hours before stimulation, the culture medium was replaced with MEM containing 0.2% fatty acid-free bovine serum albumin (BSA) for serum starvation. Cells were subsequently stimulated with recombinant human transforming growth factor-β1 (TGF- β1; R&D Systems, MN, USA) at a final concentration of 20 ng/mL for the indicated times. For time-course experiments, cells were stimulated with TGF- β1 for 1, 3, 6, 12, or 24 h. SIS3 (15945; Cayman Chemical Company, Ann Arbor, MI, USA), a selective inhibitor of SMAD3 phosphorylation, or U0126 (211-01051; Wako Pure Chemical Industries, Osaka, Japan), a MEK inhibitor, was added at final concentrations of 5 μM and 20 μM for 1 h and 2 h before TGF- β1 stimulation, respectively.

### RNA extraction and quantitative real-time PCR

Total RNA was extracted from skin samples obtained from patients with SSc and from healthy controls, as well as from cultured human dermal fibroblasts, using the RNeasy Fibrous Tissue Mini Kit or the RNeasy Mini Kit (Qiagen, Valencia, CA, USA), according to the manufacturer’s instructions. Complementary DNA (cDNA) was synthesized from the extracted RNA using ReverTra Ace qPCR RT Master Mix (TOYOBO, Osaka, Japan). Quantitative real-time PCR (qPCR) was performed using THUNDERBIRD SYBR qPCR Mix (TOYOBO) and an ABI Prism 7000 Sequence Detection System (Applied Biosystems, Foster City, CA, USA). The mRNA expression levels of *IER3*, *CDH2*, *COL1A1*, *COL22A1*, *BBC3*, *BAX*, *BCL2*, *CASP3*, and *CASP9* were determined.

*GAPDH* was used as the endogenous reference gene, and relative gene expression was calculated using the comparative ΔΔCT method. Primer sequences used in this study are listed in Supp. Table 1.

### Immunohistochemistry

Immunohistochemical staining for IER3 was performed using lesional skin samples from five patients with SSc and three patients with hypertrophic scars/keloids. Normal control skin samples were obtained from the uninvolved margins of benign skin tumors excised from three healthy individuals. Section staining was performed as previously described [18]. Briefly, paraffin-embedded tissue sections (3 μm thick) were deparaffinized and subjected to heat-induced antigen retrieval in Tris-EDTA retrieval buffer (pH 9.0) using a microwave at 95°C for 5 min. After blocking, sections were incubated overnight at 4°C with rabbit anti-IEX-1/IER3 polyclonal antibody (1:200; ab65152; Abcam, Cambridge, UK). After endogenous peroxidase activity was blocked, sections were incubated with a biotinylated goat anti-rabbit secondary antibody (Vector Laboratories, Burlingame, CA, USA), followed by incubation with the VECTASTAIN ABC Kit (Vector Laboratories) for 30 min. Immunoreactivity was visualized using 3,3’-diaminobenzidine (DAB), and sections were counterstained with Mayer’s hematoxylin.

### Small interfering RNA transfection

For small interfering RNA (siRNA)-mediated gene silencing, human dermal fibroblasts were transfected with Silencer Select Negative Control No.1 siRNA (4390843, Thermo Fisher Scientific, MA, USA), siRNAs targeting human *IER3* (Silencer Select siRNA, s16940 and s16941; Thermo Fisher Scientific), or siRNA targeting human *CDH2* (Silencer Select siRNA, s2771; Thermo Fisher Scientific) using Lipofectamine 3000 Transfection Reagent (L3000001, Thermo Fisher Scientific). siRNAs were mixed with transfection reagent in Opti-MEM (31985-047; Life Technologies, Carlsbad, CA, USA) and incubated for 15 min at room temperature before addition to the cells. The final siRNA concentration was 20 nM. Transfection efficiency was assessed by qPCR and flow cytometry analysis 48 hours post-transfection. Cells were transfected for 48 h and subsequently treated with vehicle or TGF-β1 (20 ng/mL) for the indicated times. Total RNA was then extracted, and gene expression was analyzed by quantitative real-time PCR as described above.

### Bulk RNA sequencing and data processing

Bulk RNA sequencing (RNA-seq) was performed by ImmunoGeneTeqs, Inc. (Noda city, Chiba, Japan). Polyadenylated RNA was captured from lysed cells using magnetic beads conjugated with poly(T) oligonucleotides containing sample-specific barcodes. Following bead collection and washing, complementary DNA (cDNA) was synthesized by reverse transcription. A common adapter sequence was added to the 5’ end of the cDNA through template switching based on the FLASH-seq method. The resulting cDNA was subjected to whole-transcriptome amplification (WTA) by PCR, followed by magnetic bead-based purification and size selection. The size distribution and concentration of the amplified cDNA were assessed using a microchip electrophoresis system (MultiNA; Shimadzu, Kyoto, Japan). For sequencing library preparation, the amplified cDNA was fragmented, followed by adapter ligation and index PCR to incorporate sequencing adapters and barcodes. The libraries were purified by magnetic bead-based size selection, and their size distribution and concentration were evaluated using MultiNA and quantitative PCR. Preliminary sequencing was performed using an Illumina iSeq system (Illumina, Inc., San Diego, CA) to determine and normalize library concentrations. The libraries were subsequently pooled and sequenced using an Illumina NovaSeq 6000 platform. The resulting sequencing reads were assigned to their respective samples based on barcode sequences and mapped to the reference genome for downstream analysis. Subsequent analyses, including the identification of differentially expressed genes (DEGs) and the generation of volcano plots and heatmaps, were performed by the authors using Python and R. Genes with an unadjusted raw *P*-value < 0.05 were considered differentially expressed. Volcano plots and heatmaps were generated to visualize differential gene expression patterns between the experimental groups.

### Apoptosis assay

To evaluate apoptosis, human dermal fibroblasts were divided into four experimental groups: (1) control scramble siRNA (si Ctrl) followed by vehicle treatment, (2) *CDH2* siRNA (si *CDH2*) followed by vehicle treatment, (3) si Ctrl followed by TGF-β1 treatment for 18 h, and (4) si *CDH2* followed by TGF-β1 treatment for the indicated times. Cells in each group were subsequently treated with 1 μM staurosporine (80050; AAT Bioquest, CA, USA) for 6 h. Cell death was assessed by Annexin V/propidium iodide (PI) staining followed by flow cytometric analysis. Briefly, cells were resuspended in 100 μL of Annexin V Binding Buffer (422201; BioLegend, San Diego, CA, USA) and incubated with 5 μL of Annexin V-FITC (640906; BioLegend). A PI working solution (0.1 mg/mL) was prepared by diluting PI stock solution (1 mg/mL; P378; Dojindo Laboratories, Kumamoto, Japan) in Annexin V Binding Buffer, and 5 μL of the working solution was added to each sample. Samples were incubated for 15 min at 25°C in the dark. After incubation, 400 μL of Annexin V Binding Buffer was added. Samples were analyzed using the CytoFLEX S flow cytometer (Beckman Coulter, Brea, CA, USA). Data were further processed using FlowJo software (v10.6.1, TreeStar, Ashland, OR, USA). Annexin V-positive cells (Annexin V⁺/PI⁻ and Annexin V⁺/PI⁺) were considered apoptotic cells.

### Statistical analysis

Statistical analyses and most data visualization were performed using GraphPad Prism version 10.10.0 (GraphPad Software, San Diego, CA, USA). Comparisons between two groups were performed using Welch’s *t*-test. Comparisons among multiple groups were performed using one-way analysis of variance (ANOVA), followed by Sidak’s post hoc test. A two-sided *P* value < 0.05 was considered statistically significant. Data are presented as the mean ± standard error of the mean (SEM), unless otherwise indicated.

## RESULTS

### IER3 expression is increased in dermal fibroblasts in fibrotic skin lesions

To investigate the expression pattern of IER3 in fibrotic skin diseases, we performed immunohistochemical analysis of skin tissue samples. IER3 signals were detected in epidermal keratinocytes across all examined skin conditions (Fig. 1A-D). In contrast, dermal fibroblasts in healthy skin exhibited only weak or no IER3 expression (Fig. 1A), whereas those in keloid and hypertrophic scar tissues showed markedly enhanced IER3 signals, with a tendency toward stronger staining in keloids than in hypertrophic scars (Fig. 1C-D). Increased IER3 expression was also observed in the dermis of patients with systemic sclerosis (SSc), although the staining intensity appeared lower than that in keloids and hypertrophic scars but higher than that in healthy skin (Fig. 1B). Given the visually observed differences in IER3 expression levels among fibrotic skin diseases, together with previous reports of higher expression of certain fibrosis-associated markers in keloid tissue than in scleroderma skin [19], these immunohistochemical findings suggest that IER3 might play a role in fibroblast activation associated with fibrotic skin diseases. To further validate these observations, we performed qPCR analysis using bulk skin tissue samples obtained from healthy individuals and patients with SSc. Because frozen skin samples from patients with keloids and hypertrophic scars were not available for molecular analysis, the qPCR analysis was limited to healthy and SSc skin samples. Consistent with the immunohistochemical findings, *IER3* expression was significantly elevated in SSc skin compared with healthy skin (Fig. 1E). Based on these observations, we next investigated the functional role of IER3 in human dermal fibroblasts.

**Figure 1.**
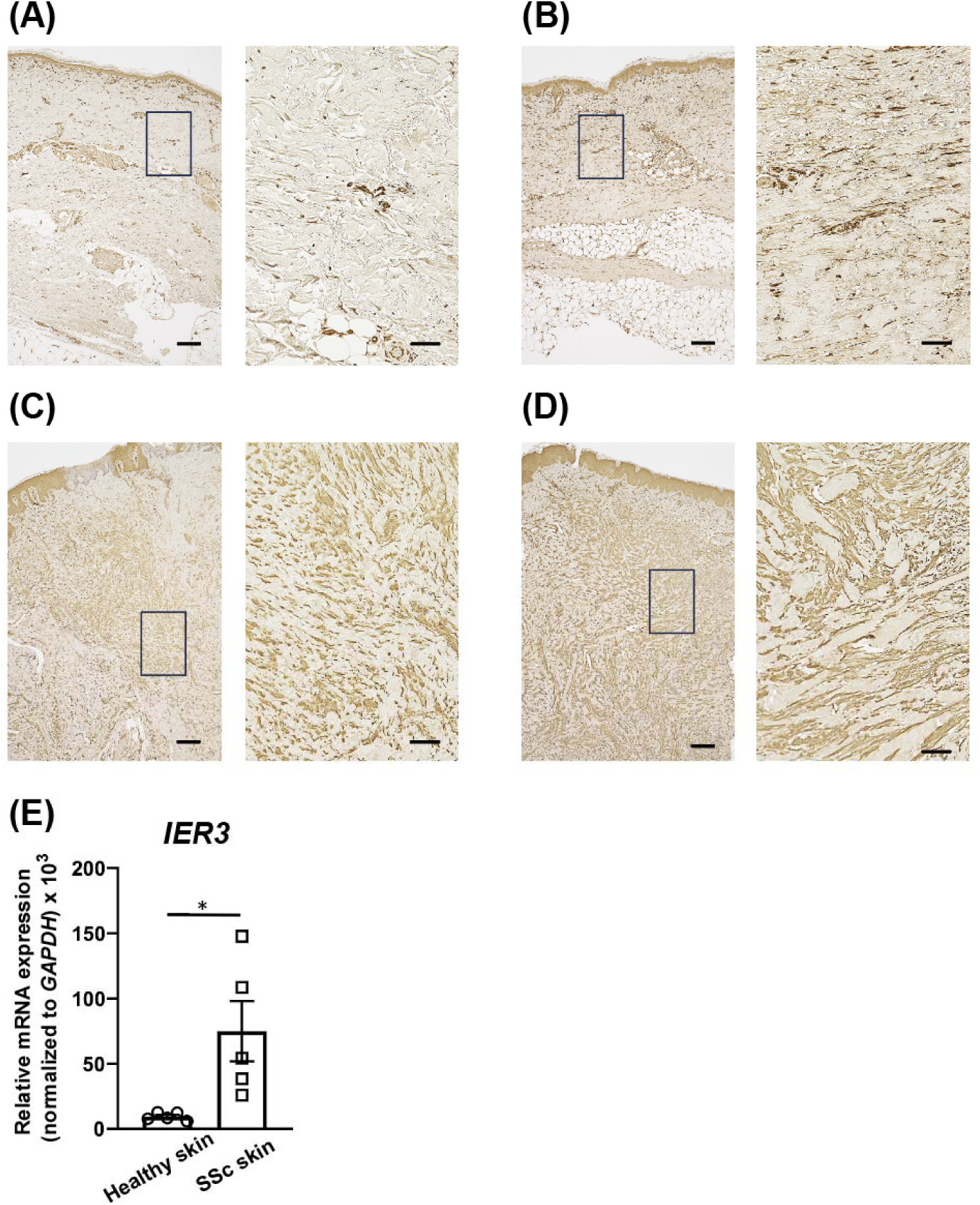
IER3 expression in fibrotic skin diseases. (A–D) Representative immunohistochemical staining of IER3 in skin sections from healthy individuals (A), patients with diffuse cutaneous systemic sclerosis (dcSSc) (B), and patients with hypertrophic scars (C) or keloids (D). Representative images were selected from skin samples obtained from three healthy individuals (n = 3), five patients with dcSSc (n = 5), and three patients with hypertrophic scars or keloids (n = 3 in total). The representative skin sections shown were obtained from the thigh of a 77-year-old healthy man (A), the right forearm of a 76-year-old woman with dcSSc and a modified Rodnan skin score (mRSS) of 39 (B), an abdominal hypertrophic scar of a 76-year-old man (C), and an abdominal keloid of a 73-year-old man (D). Higher-magnification images of the boxed areas in each panel are shown on the right. Scale bars, 200 μm in the low-magnification images and 50 μm in the high-magnification images. (E) *IER3* mRNA expression levels in skin tissue samples obtained from patients with SSc (n = 5) and healthy individuals (n = 5), as measured by quantitative real-time PCR. Statistical significance in (E) was determined using Welch’s *t*-test. *P < 0.05.

### TGF-β induces *IER3* expression in primary cultured dermal fibroblasts

Given the increased expression of IER3 in fibroblasts within fibrotic lesions, we next examined the mechanisms regulating IER3 expression in dermal fibroblasts. Primary human dermal fibroblasts isolated from healthy skin were stimulated with TGF-β1, a key profibrotic cytokine. TGF-β1 stimulation significantly increased *IER3* mRNA expression (Fig. 2A), with expression levels peaking at approximately 12-24 h after stimulation (Fig. 2B), suggesting that TGF-β1 signaling contributes to the upregulation of IER3 observed in fibrotic skin diseases. A similar induction of *IER3* expression by TGF-β1 was also observed in dermal fibroblasts derived from patients with SSc (Fig. S1A). Given previous reports that *IER3* expression is induced by hydrogen peroxide (H₂O₂), a representative inducer of oxidative stress [20], and by lysophosphatidic acid (LPA), a bioactive lipid implicated in fibrosis, in intestinal tissues [21], we next examined whether these stimuli also regulate IER3 expression in dermal fibroblasts. H₂O₂ stimulation increased *IER3* mRNA expression (Fig. 2C), whereas 18:1 LPA, a major endogenous LPA species, failed to induce *IER3* expression (Fig. S1B). To further investigate the signaling mechanisms underlying TGF-β1-induced *IER3* expression, we examined the involvement of the SMAD3 and MAPK/ERK signaling pathways, which represent canonical and non-canonical downstream pathways of TGF-β signaling, respectively [22]. TGF-β1-induced *IER3* expression was not attenuated by pretreatment with either SIS3, a selective inhibitor of SMAD3 phosphorylation (Fig. 2D), or U0126, a MEK inhibitor that suppresses MAPK/ERK signaling (Fig. 2E), although SIS3 and U0126 suppressed the TGF-β1-induced upregulation of *COL1A1* (Fig. 2D) and *COL22A1* (Fig. 2E), respectively, which were used as positive control genes based on previous studies [23, 24]. These results suggest that TGF-β-induced *IER3* expression is regulated through a mechanism independent of both MAPK/ERK signaling and SMAD3 phosphorylation.

**Figure 2.**
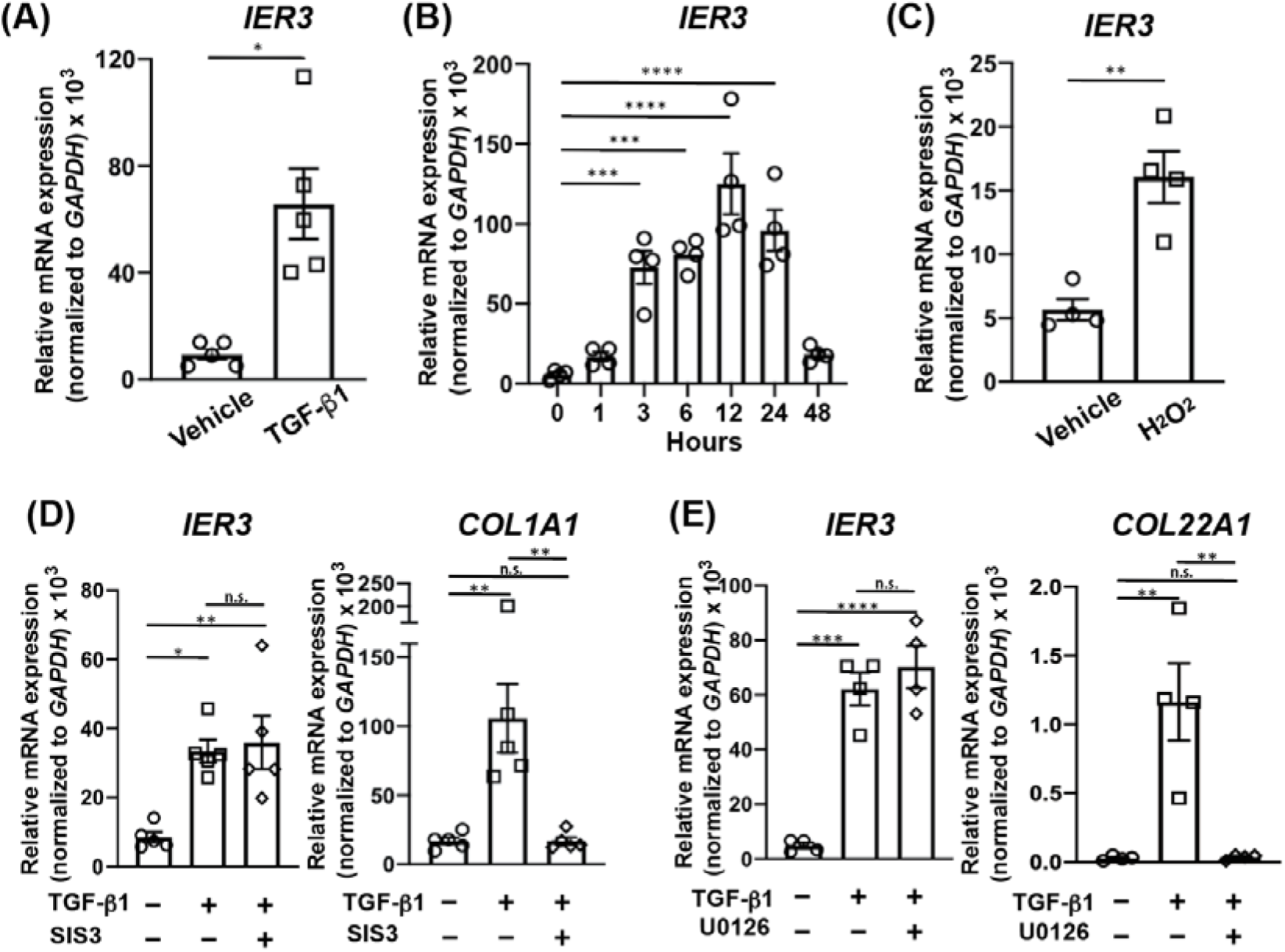
Regulation of IER3 expression by TGF-β in human dermal fibroblasts. (A) *IER3* mRNA expression levels in primary human dermal fibroblasts derived from healthy skin treated with vehicle or TGF-β1 (20 ng/mL) for 18 h following 4 h of serum starvation. *IER3* mRNA expression levels were measured by quantitative real-time PCR (qPCR) (n = 5 per group). (B) Time-course analysis of *IER3* mRNA expression in primary human dermal fibroblasts derived from healthy skin. Following 4 h of serum starvation, cells were stimulated with TGF-β1 (20 ng/mL), and *IER3* mRNA expression levels were measured by qPCR at 0, 1, 3, 6, 12, 24, and 48 h after stimulation (n = 4 per time point). (C) *IER3* mRNA expression levels in primary human dermal fibroblasts treated with vehicle or H_₂_O_₂_ (150 μM) for 2 h. IER3 mRNA expression levels were measured by qPCR (n = 4 per group). (D) *IER3* (left panel) and *COL1A1* (right panel, positive control) mRNA expression levels in primary human dermal fibroblasts treated with vehicle, TGF-β1 (20 ng/mL) alone, or TGF-β1 following pretreatment with SIS3 (5 μM), a selective inhibitor of SMAD3 phosphorylation. Cells were serum-starved for 4 h before TGF-β1 stimulation. mRNA expression levels were measured by qPCR 18 h after TGF-β1 stimulation (n = 5 per group). (E) *IER3* (left panel) and *COL22A1* (right panel, positive control) mRNA expression levels in primary human dermal fibroblasts derived from healthy skin treated with vehicle, TGF-β1 (20 ng/mL) alone, or TGF-β1 following pretreatment with U0126 (20 μM), a MEK inhibitor that suppresses MAPK/ERK signaling. Cells were serum-starved for 4 h before TGF-β1 stimulation. Cells were pretreated with U0126 for 2 h before TGF-β1 stimulation. mRNA expression levels were measured by qPCR 18 h after TGF-β1 stimulation (n = 4 for the vehicle and TGF-β1 plus U0126 groups; n = 5 for the TGF-β1 group). All mRNA expression levels were normalized to *GAPDH* expression. Each experiment was repeated at least twice, and representative data from one independent experiment are shown. Data are presented as mean ± SEM. Statistical significance was determined using Welch’s *t*-test for comparisons between two groups and one-way analysis of variance (ANOVA) followed by Sidak’s multiple comparisons test for comparisons among three groups. n.s., not significant; \**P* < 0.05, \*\**P* < 0.01, \*\*\**P* < 0.001, \*\*\*\**P* < 0.0001.

### IER3 acts as an upstream regulator of N-cadherin in TGF-β-stimulated cultured skin fibroblasts

To further elucidate the biological/functional significance of IER3 upregulation in dermal fibroblasts, we performed RNA sequencing (RNA-seq) analysis to comprehensively characterize changes in gene expression following siRNA-mediated *IER3* knockdown. The analysis was conducted under TGF-β1-stimulated conditions, in which *IER3* expression was increased (Fig. 3A). Among the genes significantly altered by *IER3* knockdown, *CDH2* was identified as one of the most prominently downregulated genes, suggesting that its expression is positively regulated by IER3 (Fig. 3A-B). *CDH2*, which encodes N-cadherin, is known to regulate cell-cell adhesion and cellular migration [25]; however, its functional role in dermal fibroblasts, particularly in fibrotic skin diseases, remains largely unexplored. We therefore focused on N-cadherin and further investigated its regulation and potential functional significance in dermal fibroblasts.

**Figure 3.**
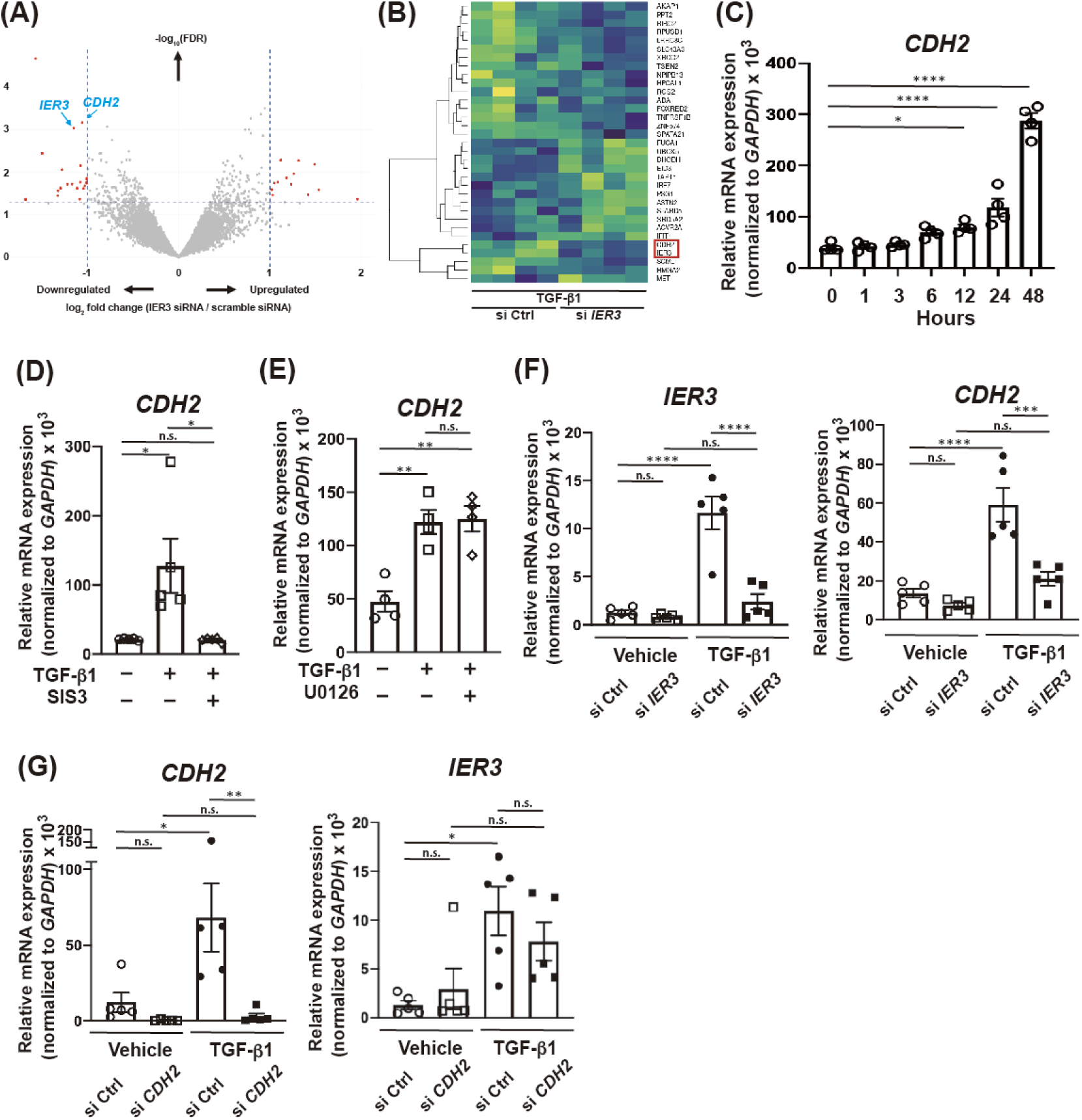
Regulation of N-cadherin expression by IER3 and TGF-β signaling in human dermal fibroblasts. (A–B) Volcano plots (A) and a heatmap (B) showing gene expression profiles obtained by RNA sequencing analysis of primary human dermal fibroblasts stimulated with TGF-β1 (20ng/ml) for 18 h following 4 h of serum starvation. Cells were transfected with scrambled siRNA (si Ctrl) or *IER3*-targeting siRNA (si *IER3*) 48 h before TGF-β1 stimulation (n = 4 per group). (C) Time-course analysis of *CDH2* mRNA expression in primary human dermal fibroblasts derived from healthy skin. Cells were serum-starved for 4 h before TGF-β1 stimulation, and *CDH2* mRNA expression levels were measured by quantitative real-time PCR (qPCR) at 0, 1, 3, 6, 12, 24, and 48 h after stimulation (n = 4 per time point). (D) *CDH2* mRNA expression levels in primary human dermal fibroblasts derived from healthy skin treated with vehicle, TGF-β1 (20 ng/mL) alone, or TGF-β1 following pretreatment with SIS3 (5 μM), a selective inhibitor of SMAD3 phosphorylation (left panel), or U0126 (20 μM), a MEK inhibitor that suppresses MAPK/ERK signaling (right panel). Cells were serum-starved for 4 h before TGF-β1 stimulation and pretreated with SIS3 or U0126 for 1 h or 2 h, respectively, before stimulation. *CDH2* mRNA expression levels were measured by qPCR 18 h after TGF-β1 stimulation (n = 5 per group). (E) Effects of *IER3* knockdown on *CDH2* mRNA expression in primary human dermal fibroblasts. Cells were transfected with scrambled siRNA (si Ctrl) or *IER3*-targeting siRNA (si *IER3*) and subsequently treated with vehicle or TGF-β1 (20 ng/mL) for 18 h, beginning 48 h after transfection and following 4 h of serum starvation. *CDH2* mRNA expression levels were measured by qPCR (n = 5 per group). (F) Effects of *CDH2* knockdown on *IER3* mRNA expression in primary human dermal fibroblasts. Cells were transfected with scrambled siRNA (si Ctrl) or *CDH2*-targeting siRNA (si *CDH2*) and subsequently treated with vehicle or TGF-β1 (20 ng/mL) for 18 h, beginning 48 h after transfection and following 4 h of serum starvation. *IER3* mRNA expression levels were measured by qPCR (n = 5 per group). All mRNA expression levels were normalized to *GAPDH* expression (C-F). Each experiment (C-G) was repeated at least twice, and representative data from one independent experiment are shown. Data are presented as mean ± SEM. (A) Differentially expressed genes were defined as those with a raw *P* value < 0.05, without correction for multiple testing (A). Statistical significance was determined using one-way analysis of variance (ANOVA) followed by Sidak’s multiple comparisons test for comparisons among three groups (C–G). n.s., not significant; *P < 0.05, **P < 0.01, ***P < 0.001, ****P < 0.0001.

We first examined the effect of TGF-β1 on *CDH2* mRNA expression in cultured dermal fibroblasts. qPCR analysis demonstrated that TGF-β1 induced *CDH2* expression in a concentration-dependent manner (Fig. 3C), with its upregulation occurring later than that of *IER3*, which was also induced by TGF-β1 (Fig. 2B). To investigate the signaling pathway responsible for this induction, cells were pretreated with either SIS3, a selective inhibitor of SMAD3 phosphorylation or U0126, a MEK inhibitor. TGF-β1-induced *CDH2* expression was markedly suppressed by SIS3 (Fig. 3D) but was not attenuated by U0126 (Fig. 3E), suggesting that *CDH2* induction by TGF-β1 is dependent on SMAD3 signaling rather than MEK/ERK signaling. In contrast, our previous findings showed that TGF-β1-induced *IER3* expression was independent of SMAD3 phosphorylation (Fig. 2C). These results indicate that, although both *IER3* and *CDH2* are induced by TGF-β1, their transcriptional regulation is mediated, at least in part, through distinct signaling mechanisms.

Consistent with the RNA-seq results (Fig. 3A-B), knockdown of *IER3* using siRNA significantly attenuated the TGF-β1-induced upregulation of *CDH2* mRNA compared with scrambled control siRNA (Fig. 3F). In contrast, although *CDH2*-specific siRNA achieved more than 90% knockdown efficiency, *CDH2* depletion did not significantly affect TGF-β1-induced *IER3* mRNA expression compared with scramble siRNA-transfected control cells (Fig. 3G). Collectively, these findings identify IER3 as an upstream regulator of N-cadherin and support the presence of a previously unrecognized unidirectional IER3-N-cadherin regulatory axis.

### CDH2 suppresses apoptosis in TGF-β-stimulated dermal fibroblasts

Next, we investigated the role of CDH2 in dermal fibroblasts using *CDH2*-specific siRNA, which achieved a knockdown efficiency of more than 95% under the TGF-β1-treatment (Fig. 3G). Given that resistance to apoptosis in activated fibroblasts and myofibroblasts is considered an important mechanism contributing to the persistence of fibrotic lesions, including keloids and hypertrophic scars [26, 27], we examined whether *CDH2* knockdown affects fibroblast susceptibility to apoptosis. Dermal fibroblasts transfected with scrambled siRNA (control) or *CDH2*-specific siRNA were treated with vehicle or TGF-β1, followed by treatment with staurosporine, which is well known to induce apoptosis [28]. Apoptosis was evaluated by flow cytometric analysis using Annexin V and propidium iodide (PI) staining (Fig. 4A). Early apoptotic cells were defined as Annexin V⁺/PI⁻, whereas late apoptotic or necrotic cells were defined as Annexin V⁺/PI⁺ [29]. Under TGF-β1-treated conditions, *CDH2* knockdown significantly increased the percentages of both early apoptotic (Annexin V⁺/PI⁻) and late apoptotic or necrotic (Annexin V⁺/PI⁺) cells compared with scrambled siRNA-transfected control cells (Fig. 4A-B). Similar trends were also observed under vehicle-treated conditions, although the differences in late apoptotic cells did not reach statistical significance (Fig. 4A-B). These findings indicate that *CDH2* knockdown enhances the susceptibility of dermal fibroblasts to staurosporine-induced apoptosis, particularly under TGF-β1 stimulation. Collectively, these findings suggest that TGF-β induces IER3 expression in dermal fibroblasts, which subsequently regulates N-cadherin expression and promotes fibroblast survival by suppressing apoptosis. This IER3–N-cadherin axis may contribute to the survival and persistence of activated fibroblasts within fibrotic skin lesions.

**Figure 4.**
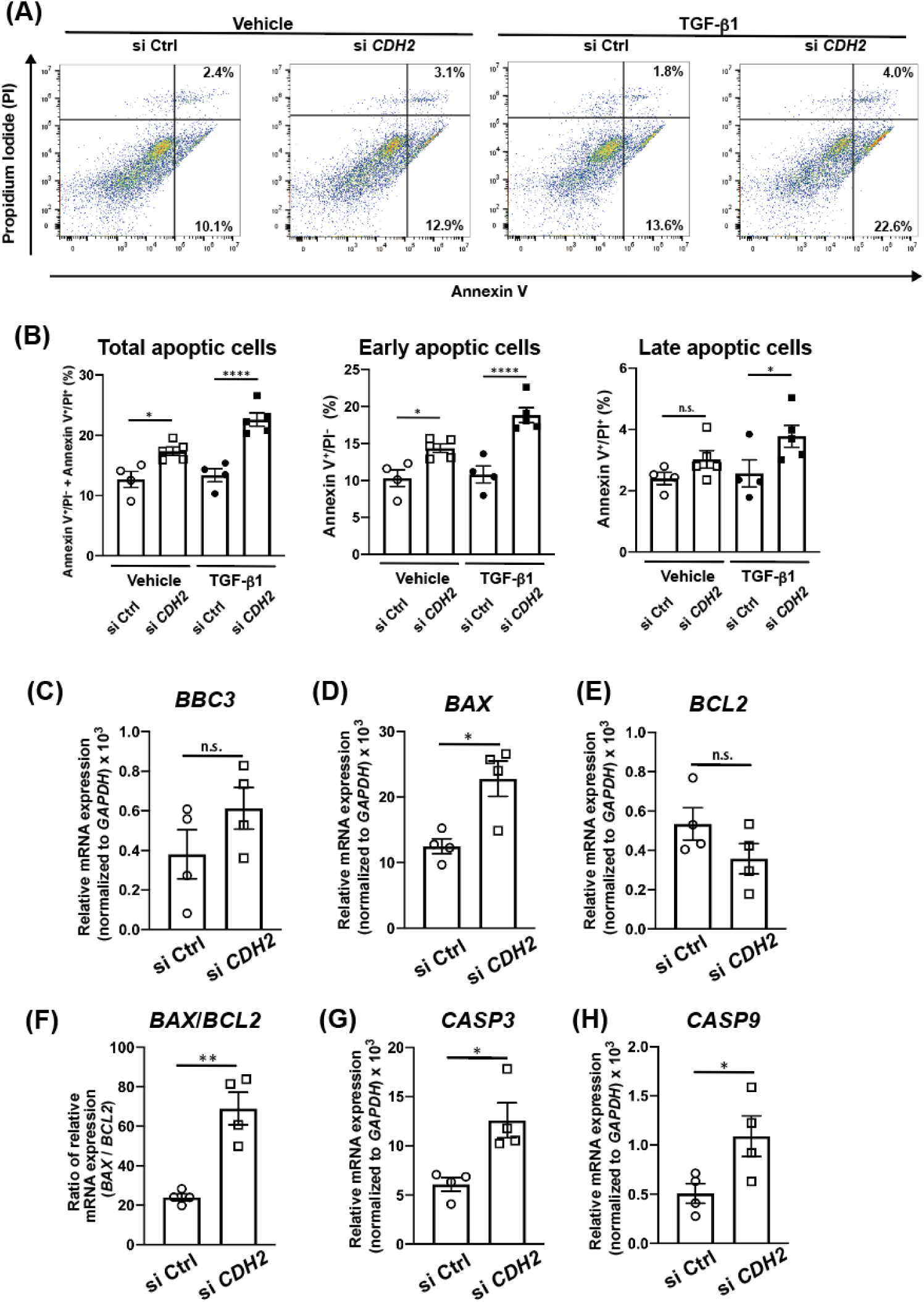
***CDH2* knockdown promotes apoptosis in human dermal fibroblasts.** (A-H) Human dermal fibroblasts were transfected with control siRNA (si Ctrl) or *CDH2*-targeting siRNA (si *CDH2*). At 48 h after transfection, cells were treated with vehicle or TGF-β1 (20 ng/mL) for 18 h, followed by stimulation with staurosporine (1 μM) for 6 h to induce apoptosis. Apoptosis was assessed by flow cytometry using Annexin V and propidium iodide (PI) staining. Representative flow cytometry plots are shown in (A). Quantification of total apoptotic cells (Annexin V⁺/PI⁻ and Annexin V⁺/PI⁺), early apoptotic cells (Annexin V⁺/PI⁻), and late apoptotic cells (Annexin V⁺/PI⁺) is shown in (B). (C-H) The mRNA expression levels of *BBC3* (C), *BAX* (D), *BCL2* (E), *CASP3* (G), and *CASP9* (H) were measured by quantitative real-time PCR and normalized to *GAPDH* expression. The *BAX*/*BCL2* expression ratio was calculated using the *GAPDH*-normalized expression levels (F). Each experiment was repeated at least twice, and representative data from one independent experiment are shown. Data are presented as mean ± SEM. Statistical significance was determined using Welch’s t-test for comparisons between two groups and one-way analysis of variance (ANOVA) followed by Sidak’s multiple comparisons test for comparisons among three or more groups. \**P* < 0.05, \*\**P* < 0.01, \*\*\*\**P* < 0.0001; n.s., not significant.

**Figure 5.**
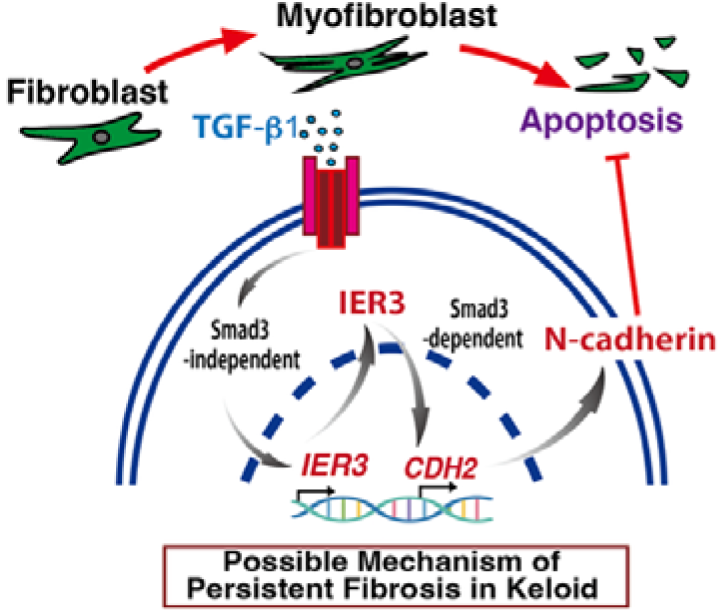
Conceptual model for the role of the IER3–N-cadherin axis in persistent fibrosis in dermal fibroblasts. TGF-β1 induces IER3 expression through a pathway independent of both SMAD3 and MAPK/ERK signaling. Subsequently, IER3 promotes N-cadherin(CDH2) expression through a SMAD3-dependent but MAPK/ERK-independent mechanism. Increased N-cadherin expression might suppress fibroblast apoptosis, thereby promoting myofibroblast persistence and contributing to sustained dermal fibrosis.

To further investigate the potential contribution of N-cadherin to apoptosis, we compared the expression of apoptosis-related genes in TGF-β1-stimulated dermal fibroblasts transfected with scrambled siRNA or *CDH2*-specific siRNA using qPCR. *CDH2* knockdown altered the expression of some apoptosis-related genes. Specifically, *BBC3* expression tended to increase, although the difference was not statistically significant (Fig. 4C). The expression of the pro-apoptotic gene *BAX* was significantly increased, whereas that of the anti-apoptotic gene *BCL2* tended to decrease without reaching statistical significance (Fig. 4D-E). Consistent with these changes, the *BAX*/*BCL2* expression ratio was increased (Fig. 4F), suggestive of a shift toward a pro-apoptotic state [30]. Furthermore, the expression levels of *CASP9* and *CASP3*, encoding an initiator and an executioner caspase, respectively, were significantly increased (Fig. 4G-H). Taken together, these findings implicate that N-cadherin may contribute to survival by suppressing mitochondria-dependent apoptosis via the intrinsic pathway in dermal fibroblasts [31].

## DISCUSSION

In the present study, we identified a previously unrecognized role of IER3 in the persistence of dermal fibroblasts under profibrotic conditions. IER3 expression was increased in dermal fibroblasts within fibrotic skin lesions, including keloids, hypertrophic scars, and SSc, and was particularly enriched in fibroblast populations in keloids and hypertrophic scars. In primary human dermal fibroblasts, TGF-β1 induced *IER3* expression, and transcriptomic analysis following *IER3* knockdown identified *CDH2* as a major downstream target of IER3. Although both *IER3* and *CDH2* were induced by TGF-β1, their regulation appeared to involve distinct signaling mechanisms: TGF-β1-induced *CDH2* expression was dependent on SMAD3 phosphorylation, whereas induction of *IER3* was not significantly affected by inhibition of either SMAD3 or MEK/ERK signaling. Importantly, *IER3* knockdown attenuated TGF-β1-induced *CDH2* expression, whereas *CDH2* knockdown did not affect *IER3* expression, indicating a unidirectional regulatory relationship between these molecules. Furthermore, *CDH2* knockdown enhanced apoptosis and increased the expression of molecules associated with the intrinsic apoptotic pathway. Collectively, these findings suggest that the IER3–N-cadherin axis represents a previously unrecognized mechanism that promotes the survival and persistence of fibroblasts in a profibrotic microenvironment.

Fibrosis is characterized not only by enhanced activation and extracellular matrix production by fibroblasts but also by the persistence of activated fibroblasts and myofibroblasts within affected tissues [1]. During normal wound healing, activated fibroblasts expand transiently and are subsequently eliminated or inactivated as tissue repair resolves. In contrast, in fibrotic skin disorders such as keloids and scars, impaired clearance of activated fibroblasts may allow sustained extracellular matrix production and thereby contribute to persistent fibrosis [3, 32]. Resistance of fibroblasts and myofibroblasts to apoptosis has therefore been proposed as an important mechanism underlying pathological fibrosis. Our findings extend this concept by identifying IER3 and its downstream molecule N-cadherin as potential regulators of fibroblast survival. The increased expression of IER3 observed in several distinct fibrotic skin disorders further suggests that this pathway may represent a common mechanism contributing to fibroblast persistence rather than a disease-specific alteration.

One of the most notable findings of the present study was the identification of N-cadherin as a downstream effector of IER3. *CDH2* encodes N-cadherin, a classical cadherin involved in cell–cell adhesion, cytoskeletal organization, and cell survival [33]. Although N-cadherin has been extensively studied in development, epithelial-to-mesenchymal transition, and cancer biology [34], its role in fibrotic fibroblasts is less well characterized. CDH2 expression has been reported to be increased in TGF-β-treated primary lung fibroblasts and in lung tissue from patients with idiopathic pulmonary fibrosis [35], and N-cadherin depletion reduces the expression of profibrotic markers including collagen I and α-SMA [35]. In addition, aberrant processing and localization of N-cadherin have been observed in pathological myofibroblasts and fibrotic tissues from multiple organs [36]. These observations, together with our findings in dermal fibroblasts, suggest that dysregulation of *CDH2* expression and/or N-cadherin function may contribute to fibroblast activation and the persistence of fibrosis across different organs.

Our results further reveal a potential mechanism through which N-cadherin may contribute to fibrosis. *CDH2* knockdown significantly increased the susceptibility of TGF-β1-treated dermal fibroblasts to staurosporine-induced apoptosis. This effect was accompanied by increased *BAX* expression and an increased *BAX*/*BCL2* ratio, as well as increased *CASP3* and *CASP9* expression, whereas *BBC3* expression was not significantly altered. These findings are consistent with enhanced activation or priming of the mitochondria-dependent intrinsic apoptotic pathway following *CDH2* depletion. N-cadherin-mediated cell adhesion can activate intracellular survival signaling in several cellular contexts, raising the possibility that increased N-cadherin expression provides activated fibroblasts with a survival advantage within fibrotic tissues. Thus, N-cadherin may actively contribute to the persistence of pathogenic fibroblasts.

Based on observations in our study, we propose a model in which sustained profibrotic stimulation induces IER3 in dermal fibroblasts, which in turn facilitates N-cadherin expression and increases resistance to apoptosis. This mechanism may be particularly important in chronic fibrotic lesions, where persistent TGF-β signaling and cellular stress coexist. Under physiological wound-healing conditions, removal of activated fibroblasts contributes to resolution of tissue repair. In pathological fibrosis, such as that observed in keloids, however, activation of the IER3–N-cadherin axis may shift this balance toward fibroblast survival, resulting in prolonged persistence of ECM-producing cells and consequently promoting maintenance of fibrotic lesions. This model provides a potential mechanistic link between cellular stress responses, TGF-β signaling, and defective fibroblast clearance.

The present findings may also have therapeutic implications. Direct inhibition of TGF-β signaling is challenging because of the pleiotropic physiological functions of TGF-β in immune regulation and tissue homeostasis. Targeting downstream mechanisms that are preferentially activated in pathogenic fibroblasts may therefore represent an alternative strategy. Our observation that IER3 expression is relatively low in fibroblasts in healthy skin but increased in fibroblasts within fibrotic lesions raises the possibility that the IER3–N-cadherin pathway could provide such a target. In particular, interference with N-cadherin-dependent survival signaling might selectively increase the susceptibility of activated fibroblasts to apoptosis and facilitate resolution of established fibrosis. Further studies will be required to determine whether pharmacological modulation of this pathway can reduce fibrosis *in vivo*.

Several limitations of this study should be acknowledged. First, the molecular mechanism by which IER3 regulates N-cadherin expression remains unclear. Because IER3 is primarily recognized as a regulator of intracellular signaling pathways rather than a classical DNA-binding transcription factor [14], intermediate signaling molecules or transcriptional regulators are likely to mediate its effects on SMAD3-dependent *CDH2* transcription. Second, although our apoptosis experiments and expression analysis of *BAX*, *BCL2*, *CASP3*, and *CASP9* support the involvement of the intrinsic apoptotic pathway, direct assessment of mitochondrial apoptosis, including mitochondrial membrane potential, cytochrome c release, and activation of caspase-9 and caspase-3 at the protein level, will be required to establish this mechanism more definitively. Third, our functional experiments were primarily performed in cultured human dermal fibroblasts. Whether inhibition of IER3 or N-cadherin promotes fibroblast clearance and ameliorates established fibrosis *in vivo* remains to be determined. Finally, because increased IER3 expression was observed in multiple fibrotic skin disorders, future studies should determine whether the IER3–N-cadherin axis is also operative in fibrosis of other organs.

In conclusion, our findings identify IER3 as a stress-responsive regulator of fibroblast persistence in fibrotic skin disease and reveal N-cadherin as an important downstream effector linking IER3 to resistance to apoptosis. We propose that persistent activation of the IER3–N-cadherin axis under profibrotic conditions promotes survival of activated fibroblasts and thereby contributes to the maintenance of pathological fibrosis. Targeting this fibroblast survival pathway may therefore represent a novel strategy for promoting the resolution of established fibrotic lesions.

### CRediT Author Statement

Ryuzo Ichimura: Validation, Formal analysis, Investigation, Data curation, Writing-review and editing, Visualization, Hibari Nakajima: Investigation, Ayaka Sugimori: Investigation, Funding acquisition, Hayakazu Sumida: Conceptualization, Methodology, Validation, Formal analysis, Investigation, Resources, Data curation, Writing–original draft preparation, Visualization, Supervision, Project administration, Funding acquisition

### Data Availability Statement

The original data analyzed in this study are available from the corresponding author upon reasonable request.

## Acknowledgements

We thank our colleagues for their help with clinical sample collection. This work was financially supported by KAKENHI grant numbers JP24K02469, JP23K18286 (to H.S.), and JP26K19379 (to A. S.) from the Japan Society for the Promotion of Science (JSPS); AMED-CREST JP23gm1210013 (to H.S.) from the Japan Agency for Medical Research and Development; grants from Maruho Takagi Dermatology Foundation (to H.S.).

