## Supplementary Figures and Tables for "IER3-N-cadherin axis contributes to apoptosis resistance in dermal fibroblasts under profibrotic conditions"

**Supplementary Information**

### Supplementary Figures

#### Figure S1

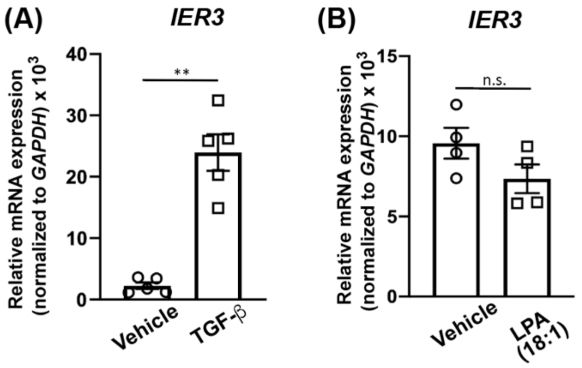


**Supplementary Figure 1.**

**Regulation of IER3 expression in human dermal fibroblasts.**

(A) *IER3* mRNA expression levels in primary human dermal fibroblasts derived from patients with systemic sclerosis (SSc) treated with vehicle or TGF-β1 (20 ng/mL) for 18 h. *IER3* mRNA expression levels were measured by quantitative real-time PCR (qPCR) (n = 5 per group). (B) *IER3* mRNA expression levels in primary human dermal fibroblasts treated with vehicle or 18:1 lysophosphatidic acid (LPA, 10 μM) for 18 h. *IER3* mRNA expression levels were measured by qPCR (n = 4 per group). Each experiment was repeated at least twice, and representative data from one independent experiment are shown. Data are presented as mean ± SEM. Statistical significance was determined using Welch's *t*-test for comparisons between two groups. n.s., not significant; $^{**}P < 0.01$.

### Supplementary Tables

#### Supplementary Table 1

**List of primers used for real‐time PCR.**

Human

| Gene | Forward Primer (5'-3') | Reverse Primer (5'-3') |
|---|---|---|
| *IER3* | CGGTCCTGAGATCTTCACCTTC | CTTTTGGCTGGGTTCGGTTC |
| *CDH2* | CCATCATTGCCATCCTGCTC | CTTCTTCTCCTCCACCTTCTTCATC |
| *COL1A1* | GCTTCACCTACAGCACCCTTGT | GATTGGGGTGGAGGGAGTTT |
| *COL22A1* | CTGTGGTGCAAAGTACTGAGGATGT | ACTCGACTGCCTTGTTTTCACCAT |
| *BBC3* | GGATGGCGGACGACCTCAAC | CCCATGATGAGATTGTACAGGACCC |
| *BAX* | ACATGGAGCTGCAGAGGATGATTG | GTTCTGATCAGTTCCGGCACCTT |
| *BCL2* | GGGGTCATGTGTGTGGAGAGC | ACAGCCAGGAGAAATCAAACAGAGG |
| *CASP3* | TGGAAGCGAATCAATGGACTCTGG | AGGTTTGCTGCATCGACATCTGT |
| *CASP9* | TTGGTGATGTCGGTGCTCTTGAG | ACTCACGGCAGAAGTTCACATTGT |
| *GAPDH* | CACCCACTCCTCCACCTTTG | CTCTCTTCCTCTTGTGCTCTTGCT |
